# GABA_B_ receptor auxiliary subunits modulate Cav2.3-mediated release from medial habenula terminals

**DOI:** 10.1101/2020.04.16.045112

**Authors:** Pradeep Bhandari, David Vandael, Diego Fernández-Fernández, Thorsten Fritzius, David Kleindienst, Jacqueline Montanaro, Martin Gassmann, Peter Jonas, Akos Kulik, Bernhard Bettler, Ryuichi Shigemoto, Peter Koppensteiner

## Abstract

The connection from medial habenula (MHb) to interpeduncular nucleus is critical for aversion- and addiction-related behaviors. This pathway is unique in selective expression of R-type voltage-gated Ca^2+^ channels (Cav2.3) in its terminals, and robust potentiation of release via presynaptic GABA_B_ receptors (GBRs). To understand the mechanism underlying this peculiar GBR effect, we examined the presynaptic localization and function of Cav2.3, GBR, and its auxiliary subunits, K^+^-channel tetramerization domain-containing (KCTD) proteins. We found selective co-expression of KCTD12b and Cav2.3 at the presynaptic active zone. GBR-mediated potentiation remained intact in KCTD12b KO mice but lasted significantly shorter. This impairment was associated with increased release and an insertion of KCTD8 into the active zone. In heterologous cells, we found direct binding of KCTD8 and KCTD12b to Cav2.3, and potentiation of Cav2.3 currents by KCTD8. The unexpected interaction of Cav2.3 with KCTDs therefore provides a means to scale synaptic strength independent of GBR activation.

## Introduction

The medial habenula (MHb) is an epithalamic structure that exclusively projects to the interpeduncular nucleus (IPN), with the dorsal MHb projecting to the lateral IPN and the ventral MHb projecting to the rostral and central subnuclei of the IPN (Figure 1A). This pathway is involved in various behaviors, including nicotine addiction and aversion (Agetsuma et al., 2010; Koppensteiner et al., 2016; Koppensteiner et al., 2017; Melani et al., 2019; Zhang et al., 2016; Zhao-Shea et al., 2013). A striking property of the MHb-IPN pathway is the prominent presynaptic localization of the R-type voltage-gated Ca^2+^ channel 2.3 (Cav2.3), a channel mainly located in postsynaptic elements in other brain areas (Parajuli et al., 2012). Furthermore, activation of presynaptic GABA_B_ receptors (GBRs) on MHb terminals exerts an unusual facilitatory effect by increasing neurotransmitter release up to ten fold (Zhang et al., 2016) and this effect appears to be involved in synaptic plasticity (Koppensteiner et al., 2017). However, this potentiation via presynaptic GBR activation only occurs in the cholinergic pathway from the ventral MHb to the rostral/central IPN, but not in the substance-Pergic pathway from the dorsal MHb to the lateral IPN (Melani et al., 2019). The presynaptic potentiation of release from ventral MHb terminals by GBR activation strongly contrasts the inhibitory effects it exerts on neurotransmitter release in other neuronal circuits (Gassmann and Bettler, 2012). The underlying mechanism is hypothesized to involve a modulation of Cav2.3 in habenular terminals via GBR-mediated signaling based on the observations that deletion of either Cav2.3 or GBRs in cholinergic MHb neurons abolished the facilitatory effect of GBR activation (Zhang et al., 2016). However, previous studies consistently reported inhibitory effects of GBRs on currents through Cav2.3 (Berecki et al., 2014; Wu et al., 1998), suggesting that a novel mechanism specific to ventral MHb terminals underlies this peculiar GBR effect. Auxiliary K^+^ channel tetramerization domain-containing (KCTD) subunits of GBRs may play a role in the GBR-mediated potentiation of neurotransmitter release, as KCTD subunits show distinct expression patterns in MHb neurons.

**Figure 1:**
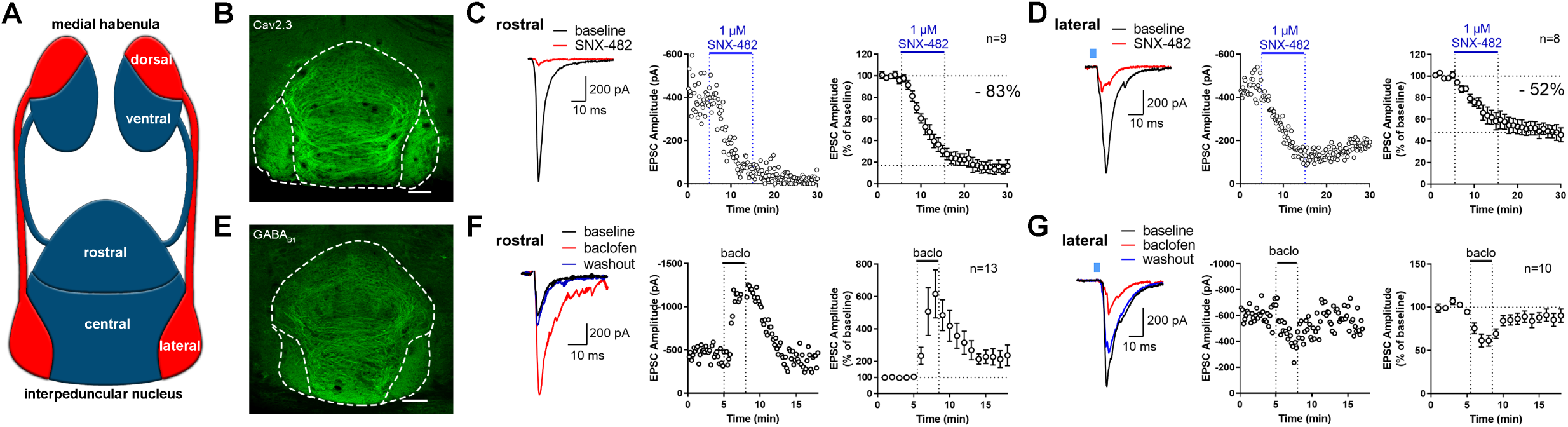
Expression and function of Cav2.3 and GABA_B_ receptors at two parallel MHb-IPN pathways. **A** Schematic drawing of the two MHb-IPN pathways. In red: the dorsal part of the MHb projects to the lateral subnuclei of the IPN. In blue: the ventral part of the MHb projects to the rostral/central subnuclei of the IPN. **B** Confocal image of Cav2.3 immunofluorescence signal indicates Cav2.3 presence in MHb axonal projections of both MHb-IPN pathways. **C** Pharmacological inhibition of Cav2.3 with SNX-482 in whole-cell recordings of rostral IPN neurons. Left: example traces before and after the application of SNX-482; middle: example time course of EPSC amplitude reduction by SNX-482; right: averaged time course of relative EPSC amplitude reduction by SNX-482. EPSC amplitudes were reduced by 83% on average (n=9 cells/9 mice). **D** In Tac1-ChR2-EYFP mice, SNX-482 reduced light-evoked glutamatergic EPSC amplitudes on average by 52% (n=8 cells/4 mice). **E** Confocal image of GABA_B1_ immunofluorescence signal indicates the presence of GABA_B_ receptors (GBRs) in all IPN subnuclei. **F** In whole-cell recordings of rostral IPN neurons, activation of GBRs by baclofen (1 µM) produced a potentiation of electrically evoked EPSC amplitudes. Left: example EPSC traces before (black) and during the application of baclofen (red) and after washout of baclofen (blue); middle: example time course of EPSC amplitudes in one cell; right: averaged time course of relative EPSC amplitude change after baclofen (n=13 cells/9 mice). **G** Baclofen reduced the amplitude of light-evoked glutamatergic EPSCs in lateral IPN neurons (n=10 cells/5 mice). Scale bars in panels (**B**) and (**E**) are 100 µm. Averaged data is presented as mean ± SEM.

There are four auxiliary KCTD subunits of GBRs, KCTD8, 12, 12b and 16. These subunits form hetero- and homo-pentamers that modulate GBR signaling kinetics (Fritzius and Bettler, 2019; Schwenk et al., 2010; Zheng et al., 2019). The expression patterns of KCTD subunits in the MHb are unique: With the exception of a weak expression in the cerebellum and superior colliculus, KCTD8 is exclusively and strongly expressed in MHb and, to a lesser extent, IPN neurons. Furthermore, KCTD12b is exclusively expressed in the ventral part of the MHb (Metz et al., 2011). In contrast, KCTD12 is weakly expressed in the ventral part of the MHb while KCTD16 is expressed in most brain areas but not the MHb. Both KCTD12 and 12b induce a rapid desensitization of GBR effector channels by uncoupling the βγ subunits (Gβγ) of the guanine nucleotide-binding protein (G-protein) from effector channels (Turecek et al., 2014). KCTD12 and 12b can form hetero-pentamers with each other but also with KCTD8 and 16 (Fritzius and Bettler, 2019). Based on proteomics studies, GBRs and their auxiliary KCTD subunits co-precipitate with release machinery proteins of the presynaptic active zone, and KCTD8 and KCTD16 were found to co-purify with presynaptic Cav2.2 Ca^2+^ channels (Müller et al., 2010; Schwenk et al., 2016). However, the functional consequences of these interactions and whether other voltage-gated Ca^2+^ channels interact with KCTDs remain unknown.

Here, we studied the nano-anatomy of Cav2.3, GBR and KCTDs and their roles in the modulation of neurotransmission from the MHb to the IPN. Our results demonstrate that Cav2.3 is located in the presynaptic active zone of habenular terminals and required for neurotransmitter release. Although we found co-localization of Cav2.3 and KCTD12b at the active zone in ventral but not dorsal MHb terminals, the GBR-mediated potentiation remained intact in the ventral MHb-to-rostral IPN pathway of KCTD12b knock-out (KO) mice. However, the potentiation returned to baseline significantly faster in KCTD12b KO mice and this impairment in GBR-mediated plasticity was KCTD8-dependent. Strikingly, we found that deletion of KCTD8 or KCTD12b reduced or increased, respectively, the probability of neurotransmitter release in this pathway. Furthermore, absence of KCTD12b led to a compensatory recruitment of KCTD8 into the active zone. In heterologous cells, we found that KCTD8 and KCTD12b but not KCTD12 bind to Cav2.3. Furthermore, Cav2.3 currents were enhanced by co-expression of KCTD8 but not KCTD12b. These results suggest that synaptic strength at the MHb-IPN pathway is scaled via GBR-independent Cav2.3 – KCTD interactions in the presynaptic active zone.

## Results

### Cav2.3 mediates neurotransmission in two distinct MHb – IPN pathways

The MHb to IPN pathway comprises two major projections (Figure 1A). The dorsal MHb projects to the lateral subnuclei of the IPN and releases glutamate and substance P whereas the ventral MHb projects to the rostral and central IPN subnuclei and co-releases glutamate and acetylcholine (Aizawa et al., 2012; Melani et al., 2019; Molas et al., 2017; Ren et al., 2011). In confocal light microscopy, both the rostral/central and lateral subnuclei show prominent Cav2.3 immunofluorescence signals (Figure 1B), in accordance with a previous study showing strong and exclusive presynaptic Cav2.3 localization in the IPN (Parajuli et al., 2012). A previous report tested the functional involvement of Cav2.3 in neurotransmission from the ventral MHb to the IPN by applying Ni^2+^ (Zhang et al., 2016), an ion known to also inhibit other Ca^2+^ channels (Lee et al., 1999). To test Cav2.3-dependency of neurotransmission in both pathways, we performed whole-cell recordings from IPN neurons in acute brain slices and applied a selective Cav2.3 blocker SNX-482 (Newcomb et al., 1998). In rostral and lateral IPN neurons, 1 µM SNX-482 strongly reduced the amplitude of excitatory postsynaptic currents (EPSCs) by 83% and 52%, respectively (Figure 1C, D). The SNX-482-mediated reduction in EPSC amplitudes was significantly higher in the ventral MHb to rostral IPN pathway compared to the dorsal MHb to lateral IPN pathway (main effect of IPN subnucleus F_1, 395_ = 213.7, P < 0.0001; two-way ANOVA). Thus, both MHb-IPN pathways rely on Cav2.3 for neurotransmitter release but show slightly different SNX-482 sensitivities.

### Differential effects of GBR activation and expression patterns of KCTDs in two distinct MHb-IPN pathways

Similarly to Cav2.3, an immunofluorescence signal for GBRs was detected in all IPN subnuclei (Figure 1E). However, GBR activation has been reported to facilitate both electrically- and optogenetically-evoked neurotransmitter release in the ventral MHb-rostral/central IPN pathway (Koppensteiner et al., 2017; Zhang et al., 2016), whereas it inhibits release in the dorsal MHb-lateral IPN (Melani et al., 2019). We confirmed that rostral IPN neurons exhibit a strong increase in EPSC amplitudes following GBR activation with 1 µM baclofen (Figure 1F), whereas EPSC amplitudes were inhibited by baclofen in lateral IPN neurons (Figure 1G). Therefore, the modulation of Cav2.3-mediated release via GBRs appears fundamentally different between the two MHb-IPN pathways. In order to better understand the differential modulation of neurotransmitter release by GBRs at the two distinct pathways, we investigated the expression of GBR auxiliary subunits, KCTDs. Immunoreactivity for KCTD8 appeared strong in all IPN subnuclei (Figure 2A), consistent with its strong expression throughout MHb and presynaptic localization in IPN. However, KCTD8 is also expressed postsynaptically in the rostral and intermediate IPN subnuclei, according to the Allen Brain Atlas. In contrast, immunoreactivity for KCTD12 and 12b was only present in the rostral and central but not the lateral IPN subnuclei (Figure 2B, C), indicating their absence in the dorsal MHb-IPN pathway. While KCTD12 immunofluorescence patterns in the rostral region suggested a mostly postsynaptic expression, KCTD12b signal showed the characteristic pattern of cholinergic MHb axons inside the rostral/central IPN, suggesting a presynaptic expression. Antibody specificity for all KCTD antibodies was confirmed using the corresponding KO animals (Figure 2A-C) while the specificity of the anti-Cav2.3 and anti-GABA_B1_ antibodies has been confirmed previously (Kulik et al., 2002; Parajuli et al., 2012).

**Figure 2:**
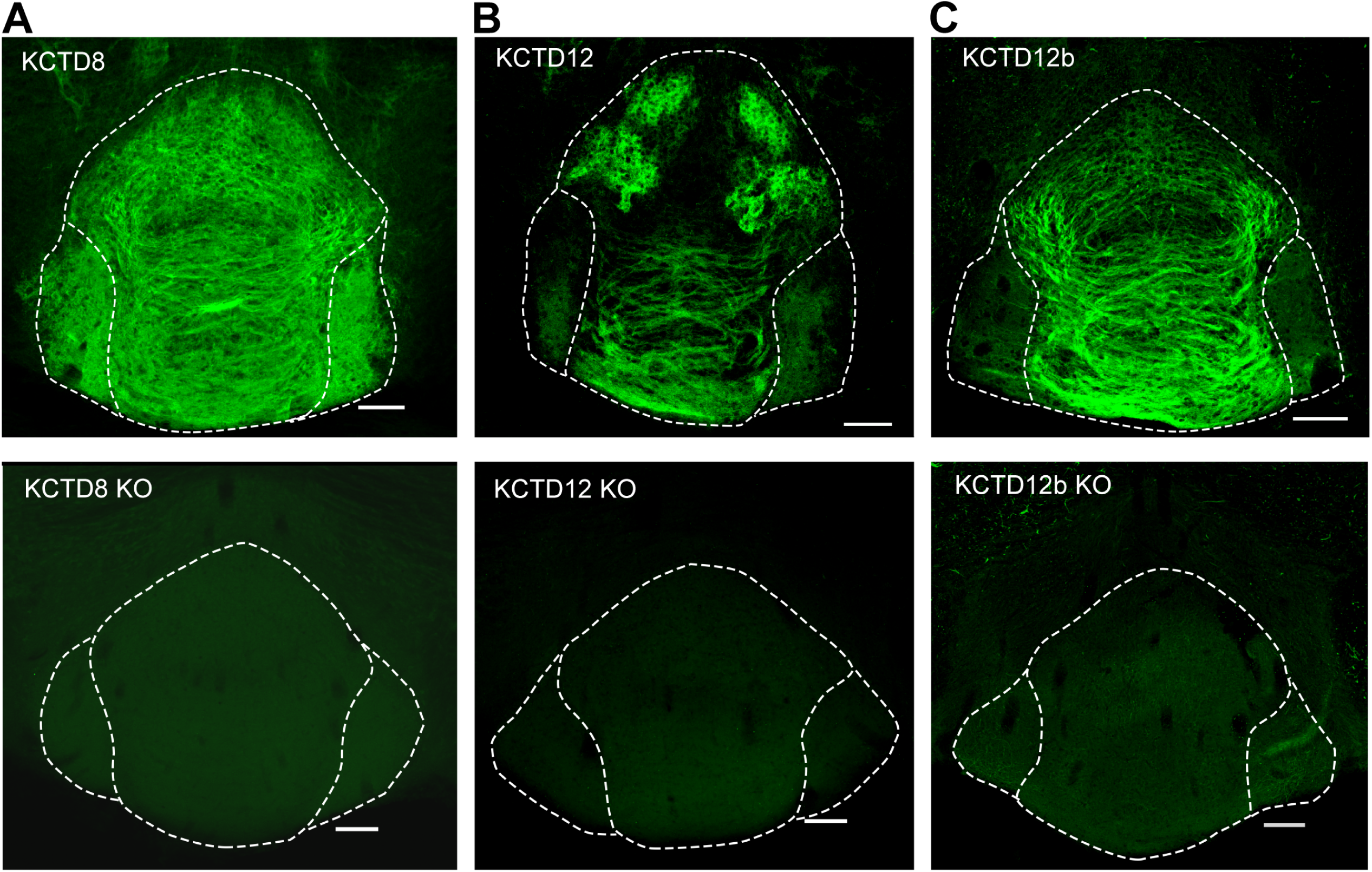
KCTDs are located in distinct IPN subnuclei. **A – C**: Confocal images of immunofluorescence signals of KCTD8, KCTD12 and KCTD12b in the IPN in WT (upper panels) and the respective KO mice (lower panels). KCTD8 immunofluorescence was present in all IPN subnuclei whereas KCTD12 and KCTD12b signals were observed only in the rostral/central but not the lateral IPN subnuclei. Scale bars: 100 µm

### Pre-embedding and freeze-fracture replica immunolabeling of presynaptic Cav2.3, GABA_B1_ and KCTDs in habenular terminals in rostral and lateral IPN subnuclei

To obtain further insight into the nanoscale distribution of presynaptic molecules, we investigated the electron microscopic localization of Cav2.3, GABA_B1_ and KCTDs in ventral and dorsal MHb terminals. We performed pre-embedding immunogold labeling and quantified the number of silver-intensified particles in the active zone, peri-synaptic region (0-50 nm distance from the edge of the active zone) and extra-synaptic regions (50-100, 100-150 and 150-200 nm distance from the edge of the active zone; Figure 3). Immunogold particles for Cav2.3 were concentrated in the active zone and, to a lesser extent, the immediate peri-synaptic region (Figure 3A). In accordance with our immunohistochemical data (Figure 2B, C), KCTD12 and KCTD12b labeling was detected only in MHb terminals in the rostral but not lateral IPN subnuclei (Figure 3D, E). In addition, particles for KCTD12 were found mostly on the postsynaptic side of asymmetrical synapses and only few synapses exhibited presynaptic KCTD12 labeling (Figure 3D, Supplementary Figure S2). In MHb terminals in the rostral IPN, Cav2.3 and KCTD12b showed similar localization patterns, with peak particle densities in the active zone and a gradual decrease in density with increased distance from the active zone (Figure 3A, E). In contrast, GABA_B1_, KCTD8 and KCTD12 showed peak localization in the peri-synaptic region with lower particle densities inside the active zone (Figure 3B – D). These results suggest that KCTD12b dominates the active zone of ventral MHb terminals in the rostral IPN whereas KCTD8 dominates the active zone of dorsal MHb terminals in the lateral IPN (Figure 3F).

**Figure 3:**
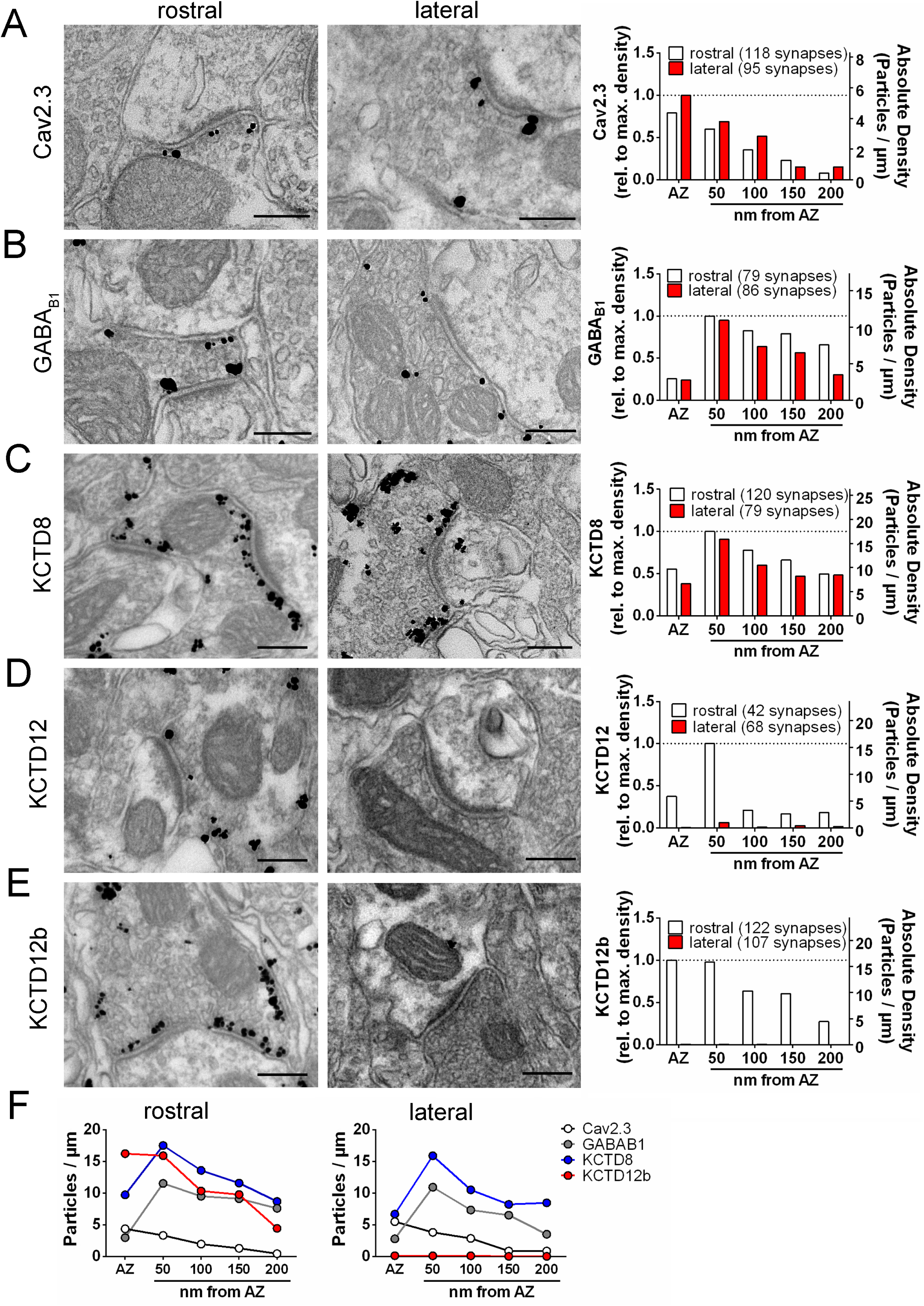
Quantification of the sub-synaptic localization of presynaptic Cav2.3, GBRs and KCTDs along both MHb-IPN pathways. Transmission electron microscopy images of 70 nm−thick sections following pre-embedding immunolabeled IPN slices for Cav2.3 (**A**), GABA_B1_ (**B**), KCTD8 (**C**), KCTD12 (**D**) and KCTD12b (**E**) from synapses in the rostral (left images) and lateral (right image) IPN subnuclei. Scale bars: 200 nm. Graph on the right displays quantification of relative and absolute silver-enhanced gold particle densities in the active zone and at distances of 50 − 200 nm from the edge of the active zone (50 nm bins). **F** Absolute labeling densities are summarized for synapses in the rostral (left panel) and lateral IPN (right panel). Note absence of KCTD12 and KCTD12b particles in presynaptic terminals inside the lateral IPN subnuclei. KCTD12 was not included in panel F because of predominantly postsynaptic localization inside the rostral IPN. Data was pooled from two animals, showing no significant difference in gold particle distribution patterns with Kolmogorov-Smirnov test (see Supplementary Figure S1).

To circumvent potential antigen-masking effects due to the protein-dense active zone region in conventional pre-embedding immunolabeling, we performed SDS-digested freeze-fracture replica labeling (SDS-FRL) (Fujimoto, 1995). This method enables unhindered access of antibodies to proteins inside/close to the pre- or postsynaptic membrane specialization and allows for multiple labeling with gold particles of distinct sizes (Indriati et al., 2013; Miki et al., 2017; Nakamura et al., 2015; Tanaka et al., 2005). In order to distinguish IPN subnuclei, we used an improved version of the grid-glued SDS-FRL method (Harada and Shigemoto, 2016) which facilitates the preservation of complete replicas during the handling procedures, a critical requirement for the identification of IPN subnuclei in the electron microscope (Figure 4A). We detected gold particles for Cav2.3 on the P-face of the presynaptic active zone and confirmed antibody specificity using Cav2.3 KO mice [Figure 4B, C, F; average density wild-type (WT): 147.8 ± 23.9 particles/µm2, n = 4 replicas; KO 3.4 ± 0.99 particles/µm2, n = 4 replicas; WT vs. KO: t_3_ = 5.87, P = 0.0099, paired t-test]. In addition, we confirmed the concentrated localization of Cav2.3 within the active zone using co-immunolabelings with a mixture of antibodies against marker proteins for the active zone, RIM1/2, neurexin and CAST (Figure 4D) (Miki et al., 2017). In the absence of marker protein labeling, demarcation of the presynaptic active zone was based on multiple criteria, including P-face curvature and intramembrane particle (IMP) size and density. The area of active zones demarcated without marker protein labeling (AZ_unmarked_) was not significantly different from that with marker labeling (AZ_unmarked_: 0.080 ± 0.0064 µm2, n = 46 AZs, AZ_marked_: 0.077 ± 0.0040 µm2, n = 80 AZs; P = 0.8692, Kolmogorov-Smirnov test) or from the area of postsynaptic IMP clusters on the E-face, the replica equivalent of the postsynaptic density seen in conventional ultrathin sections (IMP cluster: 0.077 ± 0.0059 µm2, n = 68 AZs; P > 0.9999, Kolmogorov-Smirnov test), verifying our criteria for active zone demarcation.

**Figure 4:**
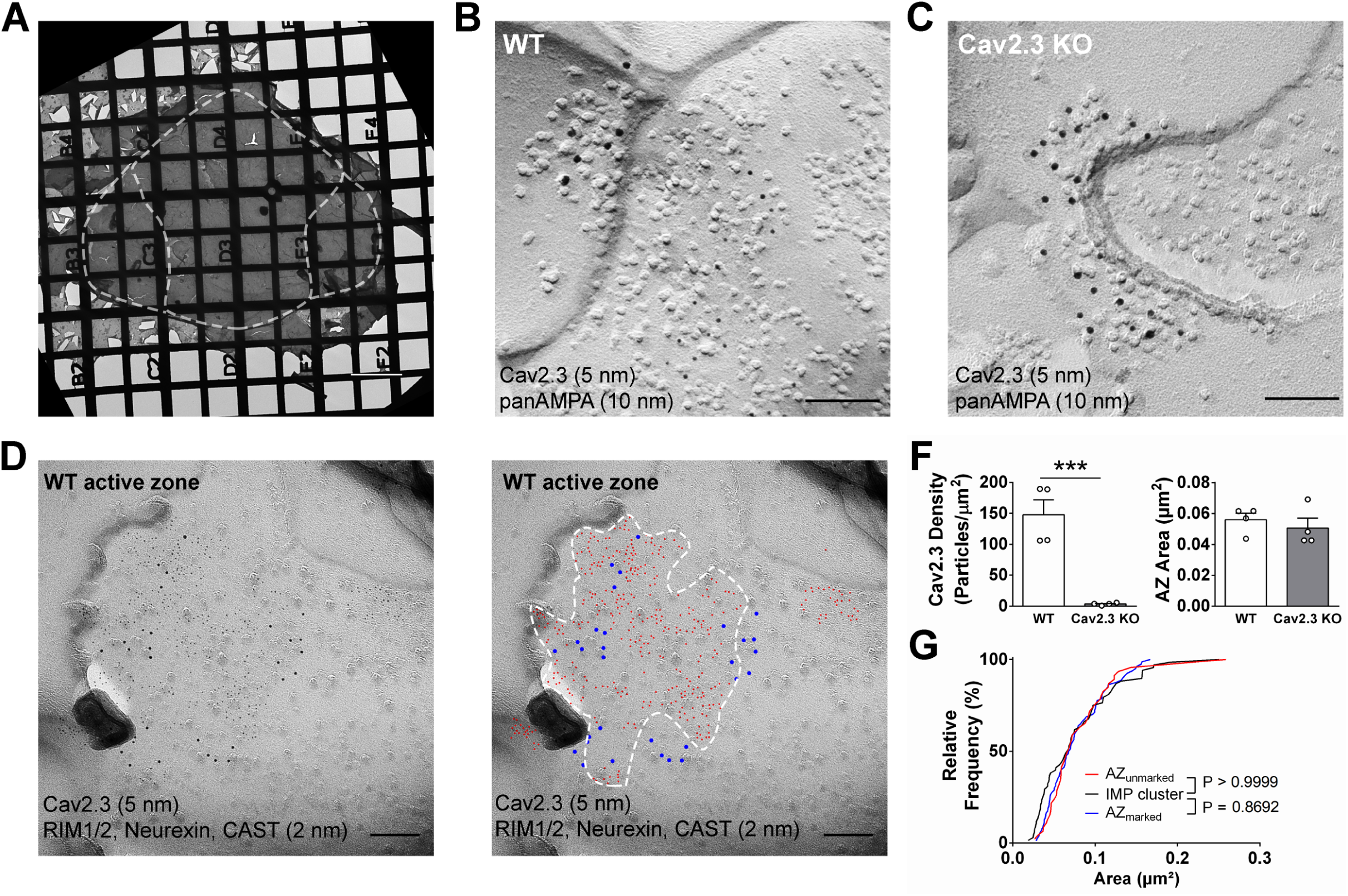
SDS-digested freeze-fracture replica labeling confirms Cav2.3 in the active zone of medial habenula terminals in the IPN. **A** Example image of a grid-glued replica containing the whole IPN. White line indicates demarcation of rostral/central and lateral subnuclei. Scale bar: 20 µm **B** Example image of a presynaptic P face and a postsynaptic E face of a habenular synapse in the rostral IPN that was double labeled with antibodies against AMPA receptors (10 nm gold) and Cav2.3 (5 nm gold). Scale bar: 100 nm. **C** Example image of a similar synaptic profile double labeled with antibodies against AMPA receptors (10 nm gold) and Cav2.3 (5 nm gold) in the rostral IPN of a Cav2.3 KO mouse. Scale bar: 100 nm. **D** Left panel: double labeling of a WT carbon-only replica with antibodies against Cav2.3 (5 nm gold) and a mixture of active zone proteins (2 nm gold), including RIM1/2, CAST and neurexin. Right panel: the same image with additional coloring of 2 nm (red) and 5 nm (blue) particles and demarcation of the active zone area based on active zone marker labeling. Scale bars: 100 nm. **F** Left panel: quantification of Cav2.3 labeling densities in the presynaptic P face in WT and Cav2.3 KO mice. *** indicates P < 0.001, unpaired t-test. Right panel: areas of demarcated active zones, including incomplete profiles, were not significantly different between replicas from WT and Cav2.3 KO mice. Data were obtained from 4 replicas from 4 mice of each genotype. **G** Comparison of active zone area demarcated with or without active zone markers and the size of the glutamatergic postsynaptic IMP clusters. P value indicates result of Kolmogorov-Smirnov test.

The result of our pre-embedding immunolabeling suggested that Cav2.3, GBR and KCTDs are localized in and around the active zone of MHb terminals (Figure 3). In order to confirm that these molecules are actually co-localized inside the same terminals, we performed double immunolabelings for Cav2.3 and either GABA_B1_, KCTD8 or KCTD12b in SDS-FRL (Figure 5). We focused on these main presynaptic KCTDs because immunogold labeling for KCTD12 was observed in few presynaptic terminals (Figure 3D) and was located mostly postsynaptically in the rostral IPN (Supplementary Figure S2). We found that in ventral MHb terminals in the rostral IPN, Cav2.3 is co-localized with GABA_B1_, KCTD8 and KCTD12b in over 97% of all Cav2.3-positive active zones (Figure 5A, B; GABA_B1_: 98 ± 0.70%, n = 3 replicas; KCTD8: 98 ± 1.10%, n = 3 replicas; KCTD12b: 97 ± 1.8%, n = 4 replicas). Similar co-localization patterns were seen in dorsal MHb terminals located inside the lateral IPN (Figure 5A, B; GABA_B1_: 99 ± 0.69%, n = 3 replicas; KCTD8: 97 ± 0.57%, n = 3 replicas), with the exception of an absence of KCTD12b. Particle numbers and densities of all tested molecules, except KCTD12b, were comparable between MHb terminals in the rostral and lateral IPN (Figure 5B). In addition, the nearest-neighbor distances (NND) of all tested molecules were significantly smaller than those obtained from simulations of randomly distributed particles (Figure 5C), suggesting that Cav2.3, GABA_B1_ and KCTDs are clustered inside the active zone.

**Figure 5:**
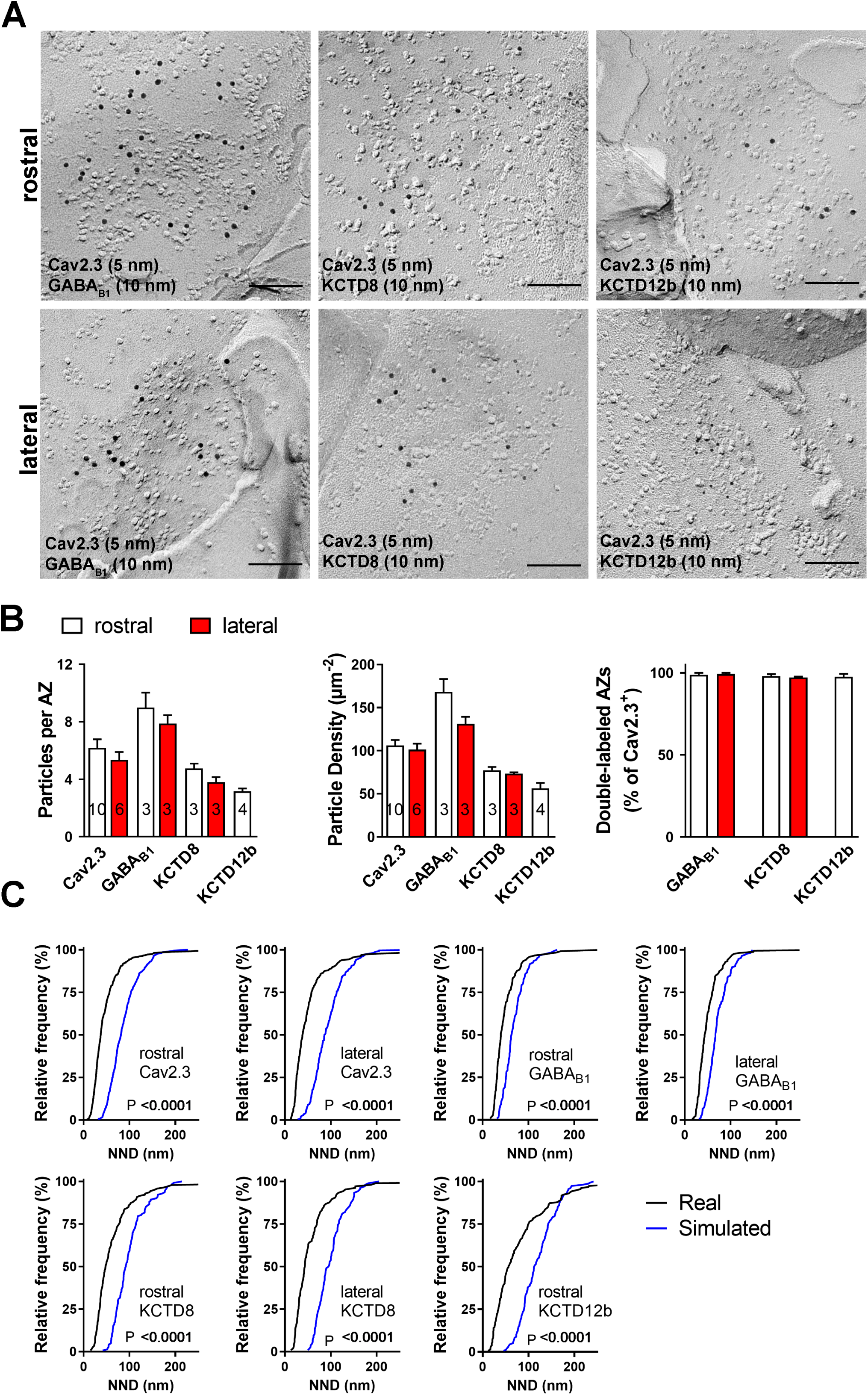
Co-localization of Cav2.3 with GBR and KCTDs in the active zone of medial habenula terminals. **A** Active zones double-labeled for Cav2.3 and either GABA_B1_ (left), KCTD8 (middle) or KCTD12b (right) in IPN replicas. Top row images are from presynaptic terminals in the rostral IPN, bottom row images are from presynaptic terminals in the lateral IPN. Scale bar: 100 nm. **B** Quantification of active zone immunolabeling in the rostral and lateral IPN. With the exception of the absence of KCTD12b in lateral IPN terminals, absolute particle numbers per active zone (left graph) and particle densities (middle graph) are comparable between MHb terminals in the rostral and lateral IPN. Right graph: Over 97% of active zones positive for Cav2.3 labeling also show labeling for one of the other molecules (GABA_B1_, KCTD8 or KCTD12b), suggesting co-localization of all presynaptic molecules inside the same active zone. Numbers inside the bars indicate the number of replicas used for each quantification. **C** Nearest neighbor distance (NND) for all presynaptic molecules in MHb terminals inside the rostral and lateral IPN based on the real (black line) and simulated random distribution (blue line). Smaller NND values in real distributions compared to simulation suggest clustering of all presynaptic molecules. P values calculated via Kolmogorov-Smirnov test.

### KCTDs modulate GBR-mediated presynaptic plasticity

Based on the selective expression of KCTD12b co-localized with Cav2.3 in the rostral/central but not lateral IPN, we tested the impact of different KCTDs on the GBR-mediated potentiation of neurotransmitter release from ventral MHb terminals in the rostral IPN (Figure 6A, B). Application of 1 µM baclofen still potentiated EPSC amplitudes in all KCTD KO lines with similar intensities (WT: 615.1 ± 148.9% of baseline; n = 13; KCTD8 KO: 470.8 ± 73.5%; n = 9; KCTD12b KO: 485.4 ± 146.6%; n = 12; KCTD8/12b double KO: 445.8 ± 76.2%; n = 10; F_4, 51_ = 0.4723, P = 0.7558, one-way ANOVA), suggesting that KCTDs are not involved in the GBR-mediated enhancement in EPSC amplitude. However, GBR-mediated potentiation was sustained significantly longer in WT and KCTD8 KO mice compared to KCTD12b KO mice (Figure 6A, B; effect of genotype: F_4, 884_ = 13.82; P < 0.0001; WT vs. KCTD12b KO: P < 0.0001; KCTD8 KO vs. KCTD12b KO: P < 0.0001; two-way ANOVA with Tukey post hoc test), suggesting an impairment of GBR-mediated plasticity in KCTD12b KO mice. Interestingly, the sustained plasticity after baclofen was re-instated in KCTD8/12b double KO mice (Figure 6B; KCTD8/12b vs. KCTD12b KO: P < 0.0001), suggesting that the effect may be mediated by KCTD8. In addition, there was no difference in the baclofen response kinetics or relative EPSC amplitude increase between WT and KCTD8 KO mice, suggesting that KCTD8 does not interfere with GBR-mediated plasticity when KCTD12b is present.

**Figure 6:**
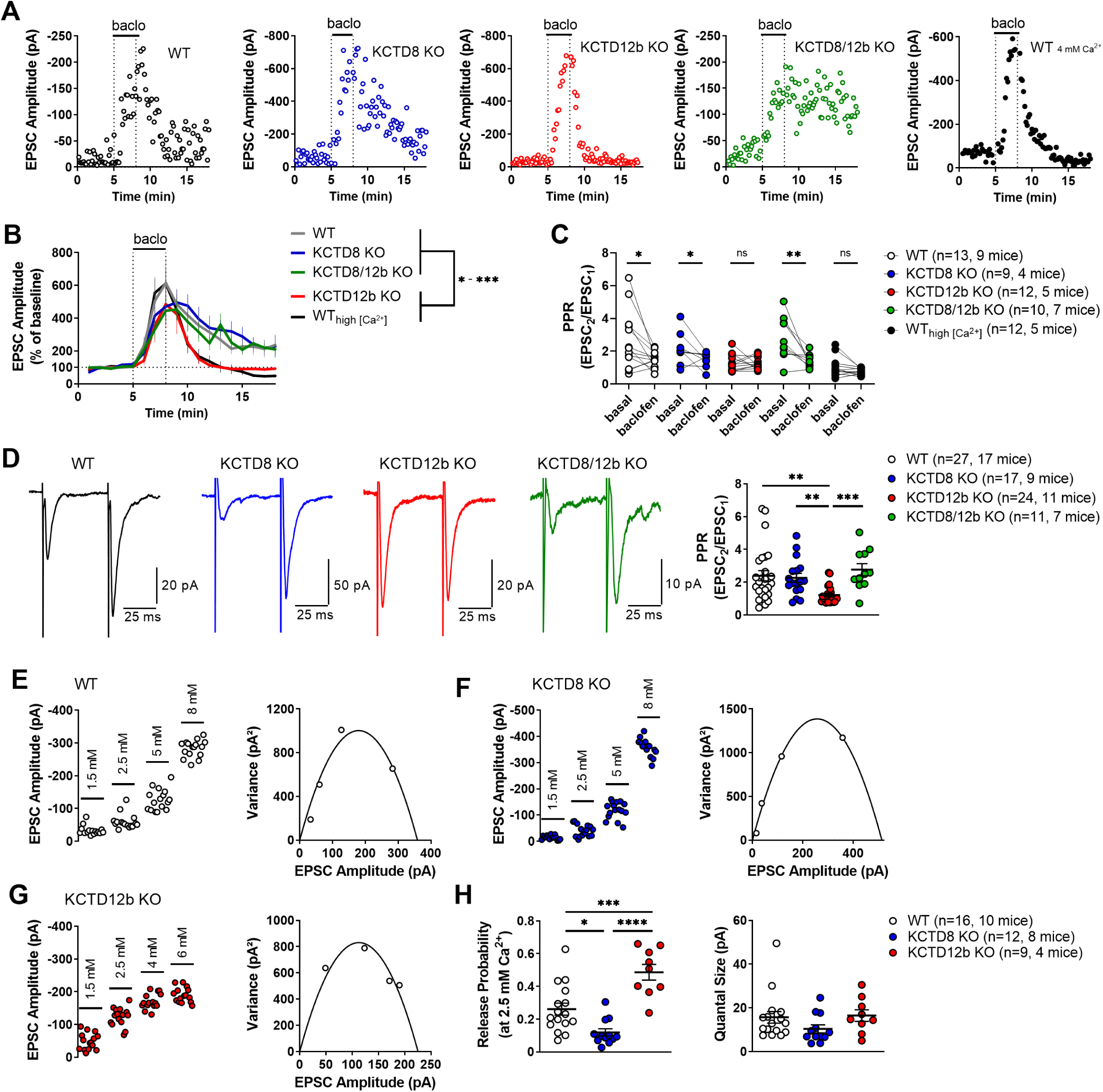
KCTD8 modulates GBR-mediated short-term plasticity by increasing basal release probability. **A** In whole-cell recordings from rostral IPN neurons, a three minute application of baclofen (1 µM) produced a strong increase in electrically evoked, glutamatergic EPSC amplitude in WT and KCTD KO mice. **B** The GBR-mediated potentiation was significantly shorter in KCTD12b KO mice and in WT mice in the presence of elevated external Ca^2+^ concentrations (4 mM) compared with all other groups, including KCTD8/12b double KO mice. Data of the WT group is identical to that presented in Figure 1E. * − *** indicates P < 0.05 − 0.001 in a two-way ANOVA with Tukey post hoc test. **C** Comparison of paired-pulse ratios (PPR) of electrically evoked glutamatergic excitatory postsynaptic currents (EPSCs) of the same neurons shown in panel (**B**) before and during the application of baclofen. Baclofen significantly reduced PPR values in WT, KCTD8 KO and KCTD8/12b double KO mice. In contrast, KCTD12b KO mice showed low basal PPR values that were not significantly affected by baclofen. PPRs of slices from WT mice in the presence of high external Ca^2+^ also remained unaffected by baclofen. * indicate P < 0.05, ** indicate P < 0.01; Wilcoxon test. **D** Basal PPR was significantly lower in KCTD12b KO mice compared with WT, KCTD8 KO and KCTD8/12b double KO mice; ** indicate P < 0.01, *** indicate P < 0.001, Kruskal-Wallis with Dunn’s post hoc test. **E-G** Example variance-mean measurements of electrically evoked EPSC amplitudes at varying external Ca^2+^ concentrations in recorded in rostral IPN neurons of WT (**E**), KCTD8 KO (**F**) and KCTD12b KO mice (**G**). **H** Quantification of release probability (left graph) and quantal size (right graph) indicates significantly reduced release probabilities in KCTD8 KO mice and increased release probabilities in KCTD12b KO mice. Quantal size remained unaffected. * P < 0.05, *** P < 0.001, **** P < 0.0001 one-way ANOVA with Tukey’s post hoc test.

### KCTDs modulate neurotransmitter release probability from ventral MHb terminals

In accordance with a previous report (Koppensteiner et al., 2017), the paired-pulse ratio (PPR) of consecutive electrically evoked EPSCs at 20 Hz in WT and KCTD8 KO mice was significantly reduced by baclofen (Figure 6C; WT basal PPR: 2.66 ± 0.49, WT baclofen PPR: 1.48 ± 0.14, n = 13 cells; P = 0.0215; KCTD8 KO basal PPR: 2.15 ± 0.32, KCTD8 KO baclofen PPR: 1.50 ± 0.15, n = 9 cells; P = 0.0391; Wilcoxon test). Surprisingly, this effect was absent in KCTD12b KO mice (Figure 6C; KCTD12b KO basal PPR: 1.30 ± 0.15, KCTD12b KO baclofen PPR: 1.25 ± 0.11, n = 12 cells, P = 0.6772, Wilcoxon test) but was reinstated in KCTD8/12b double KO mice (Figure 6D: KCTD8/12b KO basal PPR: 2.79 ± 0.41; KCTD8/12b KO baclofen PPR: 1.40 ± 0.13, n = 10 cells; P = 0.0059, Wilcoxon test). Further basal PPR recordings (Figure 6D) revealed that PPR was significantly lower in KCTD12b KO mice (1.22 ± 0.11, n = 24 cells) compared with WT (2.40 ± 0.31, n = 27 cells; P = 0.0051, Kruskal-Wallis test), KCTD8 KO (2.26 ± 0.27, n = 17 cells; P = 0.0080) and KCTD8/12b KO mice (2.76 ± 0.37, n = 11 cells; P = 0.0010). PPR values are generally thought to be inversely correlated with release probability (Dobrunz and Stevens, 1997), therefore, these findings suggest that basal release in KCTD12b KO mice is higher than in WT and KCTD8 KO mice, and that this effect is KCTD8-dependent.

To test whether the alteration in the baclofen potentiation time course in KCTD12b KO mice depends on release probability, we next tested the effect of baclofen in WT mice in the presence of high concentrations of external Ca^2+^ (4 mM instead of 2.5 mM; Figure 6A, B). Interestingly, high external Ca^2+^ concentrations led to baclofen responses that terminated similarly as those observed in KCTD12b KO mice (main effect of genotype/treatment: F_4, 884_ = 13.82; P < 0.0001; 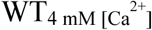 vs.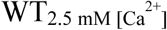: P < 0.0001; vs. KCTD8 KO: P < 0.0001; vs. KCTD12b KO: P = 0.9104; vs. KCTD8/12b KO: P = 0.0135; two-way ANOVA with Tukey post hoc test). These results suggest that the activation of presynaptic GBRs on ventral MHb terminals induces plasticity which is occluded under conditions of high basal release.

To confirm the increase in release probability in KCTD12b KO mice, we performed variance-mean analysis (Figure 6E-G) to estimate the values of release probability (at 2.5 mM external Ca^2+^) and quantal size (Figure 6H). Compared with WT mice (WT release probability: 0.26 ± 0.04, n = 16 cells), the release probability from ventral MHb terminals of KCTD12b KO mice (0.49 ± 0.05, n = 9 cells) was significantly increased (Figure 6H; F_2, 34_ = 21.23, P < 0.0001; WT vs. KCTD12b KO: P = 0.0005, one-way ANOVA with Tukey post hoc test). Surprisingly, the release probability in KCTD8 KO mice (0.12 ± 0.02, n = 12 cells) was significantly lower than that of WT or KCTD12b KO mice (WT vs. KCTD8 KO: P = 0.0176; KCTD8 KO vs. KCTD12b KO: P < 0.0001; Tukey post hoc test). The quantal size remained unaffected by genotype (F_2, 34_ = 1.613, P = 0.2142). This result suggests that KCTDs differentially modulate the probability of basal neurotransmitter release from ventral MHb terminals.

### Absence of KCTD12b induces compensatory increase in KCTD8 in the active zone of ventral MHb terminals

Next, we investigated potential nano-anatomical changes associated with the increased release probability in KCTD12b KO mice. To this aim, we performed SDS-FRL and co-immunolabeled Cav2.3/GABA_B1_ or Cav2.3/KCTD8 in replicas of WT and KCTD12b KO IPN samples (Figure 7A). Since KCTD12b is not expressed in MHb terminals in the lateral IPN, no effects on expression of these molecules are expected in the lateral IPN in KCTD12b KO mice. Thus, we normalized the densities of presynaptic molecules in the rostral IPN to the average density of the same molecule in the corresponding lateral IPN in the same replicas. Thereby, we avoided interference by technical variabilities in labeling efficiencies between individual replicas. We found no significant difference in the densities of Cav2.3 in the presynaptic active zone in the rostral relative to the lateral IPN (Figure 7B; WT rostral: 1.01 ± 0.04 fold of lateral IPN, n = 8 replicas from 8 mice; KCTD12b KO: 1.06 ± 0.13 fold of lateral IPN, n = 8 replicas from 8 mice; P = 0.5054, Mann-Whitney test). Interestingly, the relative density of KCTD8 in the active zone of MHb terminals was increased approximately 2-fold in KCTD12b KO mice compared with those of WT mice (Figure 7B; WT rostral: 0.84 ± 0.12 fold of lateral IPN, n = 5 replicas from 5 mice; KCTD12b KO: 2.09 ± 0.45 fold of lateral IPN, n = 5 replicas from 5 mice; P = 0.0079, Mann-Whitney test). This result indicates that the increased release probability in KCTD12b KO mice could be ascribable to a compensatory increase of KCTD8 in the active zone resulting in enhanced association of KCTD8 with Cav2.3. On the other hand, GABA_B1_ expression in the presynaptic active zone of MHb terminals was not significantly different between WT and KCTD12b KO (Figure 7B; WT rostral: 1.30 ± 0.18 fold of lateral IPN, n = 3 replicas from 3 mice; KCTD12b KO: 0.89 ± 0.07 fold of lateral IPN, n = 3 replicas from 3 mice; P = 0.4000, Mann-Whitney test). These results suggest that KCTDs in the active zone dynamically regulate the release probability, which may be important in scaling of synaptic strength independent from GBR effects in the MHb-IPN pathway.

**Figure 7:**
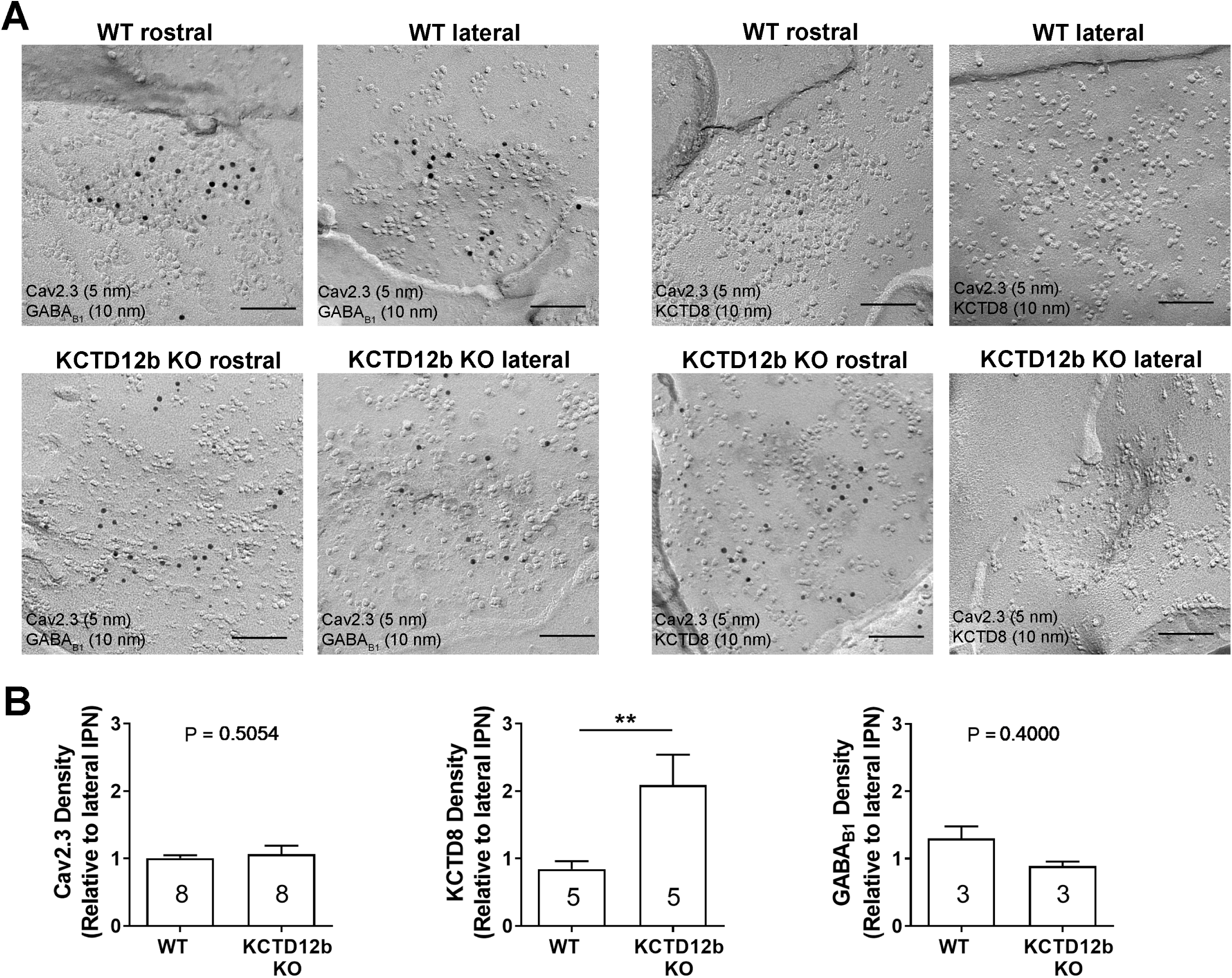
Absence of KCTD12b leads to a compensatory increase of KCTD8 inside the active zone of ventral MHb terminals. **A** Example images of active zones containing Cav2.3 and either GABA_B1_ (left panels) or KCTD8 (right panels) in replicas of WT (upper row) and KCTD12b KO IPN tissue (lower row). Scale bars: 100 nm **B** Quantification of relative densities for Cav2.3, KCTD8 and GABA_B1_ in active zones located in the rostral IPN of WT and KCTD12b KO mice. Densities were normalized to the average density in MHb terminals inside the lateral IPN of the same replica. The number inside the bars indicate the number of replicas used for quantification. ** indicate P < 0.01 in a Mann-Whitney test.

### KCTD8 and KCTD12b directly bind Cav2.3 and KCTD8 enhances currents through Cav2.3

Previous studies positioned presynaptic Ca^2+^ channels at the center of large, macromolecular complexes that also contained GBRs and KCTDs (Müller et al., 2010), and some KCTDs were found to co-purify with Cav2.2 subunit of N-type Ca^2+^ channels in the absence of GBRs (Schwenk et al., 2016). To test whether presynaptic KCTDs in MHb terminals may directly interact with Cav2.3, we performed a co-immunoprecipitation experiment in HEK293 cells transiently expressing Cav2.3 and either KCTD8, KCTD12 or KCTD12b. Interestingly, we found selective binding of Cav2.3 to KCTD8 and KCTD12b but not to KCTD12 (Figure 8A). These results suggest that KCTD8 and KCTD12b in MHb-derived axon terminals may potentially directly interact with presynaptic Cav2.3, even in the absence of GBRs.

**Figure 8:**
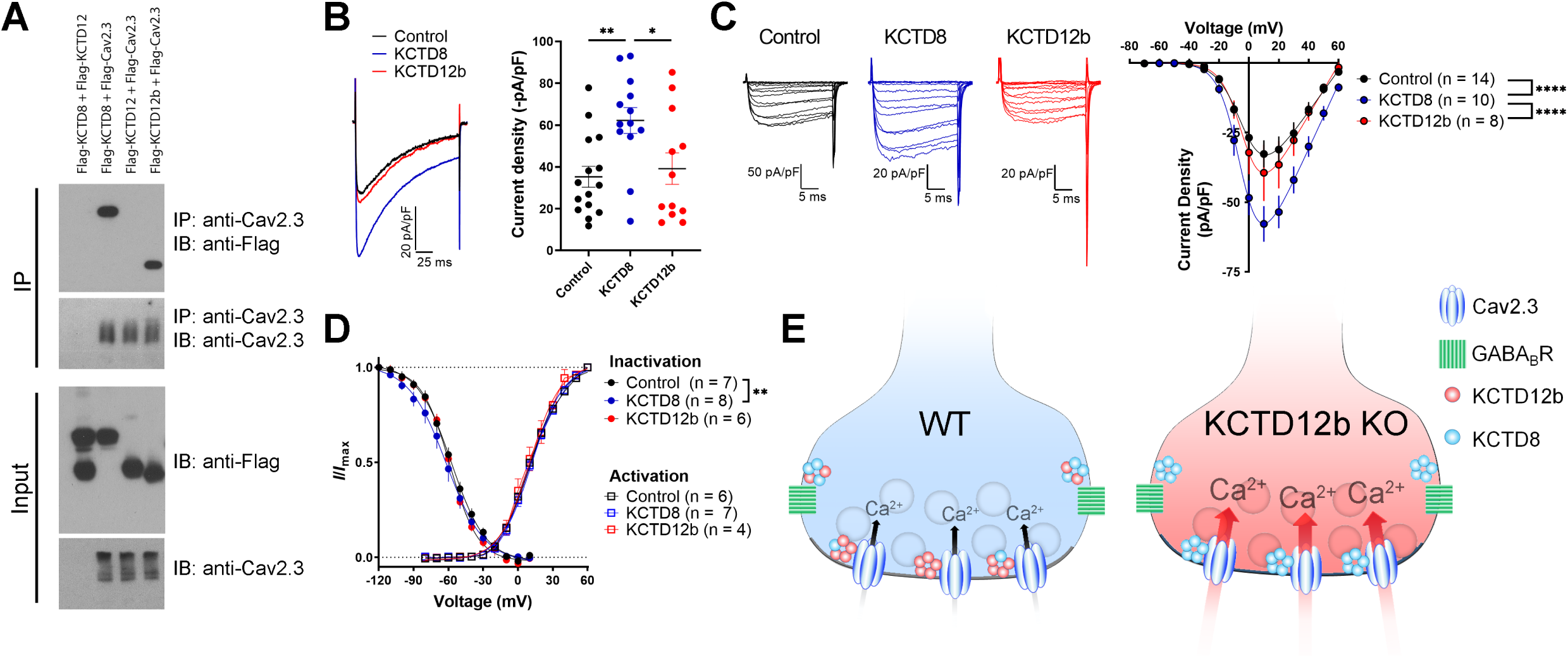
KCTDs directly bind Cav2.3 *in vitro* and KCTD8 enhances currents through Cav2.3. **A** Co-immunoprecipitation from total cell lysates of HEK293 cells transfected with Flag-tagged KCTDs and Cav2.3. Immunoprecipitation of Cav2.3 co-precipitated KCTD8 and KCTD12b, but not KCTD12. Input lanes (bottom) indicate expression of the tagged proteins in the cell lysates. **B** Whole-cell recordings from HEK293 cells stably expressing Cav2.3. Ba^2+^ current densities measured in response to a single depolarizing voltage step from −80 to 10 mV were significantly increased in KCTD8 co-transfected cells. * P < 0.05, ** P < 0.01 one-way ANOVA with Tukey post hoc test. **C** Current density-to-voltage relationship demonstrating higher current densities in KCTD8-transfected cells compared with Control- and KCTD12b-transfected cells. **** P < 0.0001 two-way ANOVA with Tukey post hoc test. **D** Activation and inactivation curves in Control-, KCTD8- and KCTD12b-transfected cells. ** P < 0.01, two-way ANOVA with Tukey post hoc test. **E** Schematic representation of the distribution and function of Cav2.3, GBRs and KCTDs in ventral MHb terminals of WT and KCTD12b KO mice. Top image: In WT terminals, the active zone contains Cav2.3 and hetero-pentameric rings comprising KCTD12b in excess over KCTD8, whereas KCTD8 and GBRs are located peri-synaptically. Bottom image: In absence of KCTD12b, KCTD8 invades the active zone and compensates for the loss of KCTD12b, resulting in increased release probability, potentially via increased Ca^2+^ influx through Cav2.3.

Using a cell line stably expressing human Cav2.3 (Dai et al., 2008), we next co-expressed either an empty vector (Control), KCTD8 or KCTD12b, and measured Ba^2+^ currents through Cav2.3. Current densities resulting from a single voltage step (–80 mV to 10 mV) were significantly increased in cells co-expressing KCTD8 compared with control and KCTD12b-transfected cells (Figure 8B, F_2, 37_ = 5.614; P = 0.0074; Control vs. KCTD8: P = 0.0086; Control vs. KCTD12b: P = 0.8959; KCTD8 vs. KCTD12b: P = 0.0382; one-way ANOVA with Tukey post hoc test). The current density-to-voltage relationship showed a significant difference between KCTD8 and both Control and KCTD12b transfected cells (Figure 8C; main effect of transfection: F_2, 406_ = 29.23, P < 0.0001; Control vs. KCTD8: P < 0.0001; Control vs. KCTD12b: P = 0.5793; KCTD8 vs. KCTD12b: P < 0.0001; two-way ANOVA with Tukey post hoc test; see also Supplementary Table 1). Furthermore, there was a slight hyperpolarization of the steady-state inactivation in the KCTD8-expressing cells compared with Control (Figure 8D; main effect of transfection: F_2, 252_ = 4.831, P = 0.0087; Control vs. KCTD8: P = 0.0059; Control vs. KCTD12b: P = 0.2667; two-way ANOVA with Tukey post hoc test) whereas the voltage-dependence of channel activation was not different between groups (Figure 8D; F_2, 201_ = 2.303; P = 0.1025, two-way ANOVA). These results suggest that the increased release probability in KCTD12b KO mice may have resulted from a KCTD8-mediated enhancement of Cav2.3 currents following the compensatory increase of KCTD8 in the active zone (Figure 8E).

## Discussion

We used a combination of anatomical, biochemical and functional methods to study the role of KCTDs at two parallel pathways from the MHb to the IPN and identified an unexpected function of KCTDs in the modulation of neurotransmitter release. We confirmed that both dorsal MHb-IPN and ventral MHb-IPN pathways require Cav2.3 for transmitter release, but only the ventral MHb-IPN pathway exhibits GBR-mediated presynaptic potentiation. KCTD8 and KCTD12b show distinct expression along these pathways, with KCTD12b specifically expressed in the cholinergic ventral MHb-IPN pathway. Using electron microscopy, we verified that Cav2.3 and KCTDs co-localize inside the presynaptic active zone of MHb terminals. Although GBR-mediated potentiation of release was still observed in KCTD KO mice, we found that loss of KCTD12b led to impaired GBR-mediated plasticity. This impairment was associated with an increase in basal release probability, suggesting occlusion of GBR-mediated plasticity due to high basal release. In contrast, absence of KCTD8 significantly reduced release probability. In addition, synapses lacking KCTD12b showed increased densities of KCTD8 in the active zone. In HEK cells, we identified direct binding of KCTD8 and KCTD12b to Cav2.3 and found a KCTD8-dependent increase in currents through Cav2.3. Thereby, our results suggest that KCTDs exert a regulatory role without GBR activation to scale synaptic strength at the MHb-IPN pathway.

### Dominant presynaptic localization and function of Cav2.3 in MHb-IPN pathway

In contrast to other brain areas, R-type Ca^2+^ channels may be the exclusive source of presynaptic Ca^2+^ influx to trigger release from MHb terminals, given that neither N-nor P/Q-type Ca^2+^ channels are expressed in MHb neurons (Ludwig et al., 1997). Although this exclusivity has been described anatomically (Parajuli et al., 2012), it was never tested functionally using the specific R-type channel blocker SNX-482. Our data confirmed that both MHb-IPN pathways heavily rely on Cav2.3 for release. Interestingly, projections from the ventral MHb to the rostral IPN were more sensitive to the peptide toxin SNX-482 than the dorsal MHb to lateral IPN pathway. However, in both pre-embedding and replica EM, we found similar Cav2.3 immunolabeling densities in the active zone of dorsal and ventral MHb terminals. Importantly, EPSC responses in the rostral IPN were evoked electrically, by placing an electrode on top of the fasciculus retroflexus while responses in the lateral IPN were evoked optogenetically in Tac1-ChR2-EYFP mice. It is conceivable that the blue light may have stimulated nearly all available ChR2^+^ terminals and reached fibers and terminals located more deeply inside the slice than the stimulating electrodes on the surface. Synapses deep inside the slice may be less accessible to peptide toxins, therefore, the actual sensitivity to SNX-482 in dorsal MHb terminals may be higher than observed. Another explanation could be that there are additional substance-Pergic inputs into the lateral IPN that derived from neurons of other brain regions which use other Ca^2+^ channels for release. However, this is unlikely, as previous studies have established that substance P signal inside the lateral IPN derives exclusively from the dorsal MHb (Contestabile et al., 1987).

### Role of KCTDs in potentiation of Cav2.3-mediated release by GBR activation

A previous report of the potentiating action of presynaptic GBRs on cholinergic MHb terminals identified Cav2.3 as a critical mediator of this effect, based on the observations that both genetic ablation of Cav2.3 as well as pharmacological inhibition of Cav2.3 with Ni^+^ prevented the potentiation (Zhang et al., 2016). Given the modulatory roles of KCTDs in shaping GBR effector responses (Fritzius et al., 2017), we initially hypothesized that the unique repertoire of KCTDs in ventral MHb neurons may be involved in the facilitatory action of GBRs. However, deletion of neither KCTD8, KCTD12b nor both prevented the potentiation of EPSC amplitudes by baclofen, suggesting a KCTD-independent effect. However, the duration of the GBR-mediated potentiation was influenced by KCTDs. It is unlikely that the impaired GBR-mediated plasticity in KCTD12b KO mice was caused by technical differences, for example due to more efficient washout of baclofen, because the experiments were performed under identical conditions for all genotypes and mice of different genotypes were recorded in a randomized manner. Furthermore, it is unlikely that the impaired short-term potentiation was due to changes in baclofen affinity because KCTDs do not affect agonist affinity (Rajalu et al., 2015). Normal GBR-mediated plasticity in KCTD8/12b double KO mice strongly suggests that the compensatory increase in KCTD8 in the active zone was responsible for the effect. Although KCTDs are known to modify deactivation kinetics of GBR effector K^+^ channels, it is unclear how an increase in KCTD8 would shorten the GBR-mediated response. In a heterologous expression system, co-expression of GBR with KCTD16, which is structurally and functionally similar to KCTD8 (Seddik et al., 2012), leads to a slowing of G-protein coupled inwardly rectifying K^+^ channel deactivation (Fritzius et al., 2017). Thus, one would rather expect a slowed termination of potentiation in the presence of more KCTD8, in case GBR-mediated potentiation results from activation of Cav2.3 through G-protein signaling.

### Regulation of Cav2.3-mediated release by KCTDs

In addition to the co-localization of Cav2.3 with KCTDs at active zone in habenular terminals, we identified a hitherto unknown direct interaction of Cav2.3 with KCTD8 and KCTD12b *in vitro*. Previous proteomics studies revealed co-precipitations of N-type/Cav2.2 Ca^2+^ channels with KCTD8 and KCTD16, but no other KCTD − Cav2 interactions were reported (Müller et al., 2010; Schwenk et al., 2016). Possibly, the use of total brain extracts may have limited detection to the most prominent protein interactions. Thus, brain-wide approaches to study presynaptic Ca^2+^ channel-interacting proteins may have failed to detect interactions uniquely occurring inside MHb terminals.

Basal levels of release probability from ventral MHb terminals were lowered below that of WT synapses by the selective KO of KCTD8. On the other hand, KO of KCTD12b resulted in significant increase of basal release probabilities. This finding of modulation of release independent of GBR activation could be explained by the current-enhancing effects of KCTD8 on Cav2.3 observed in a heterologous expression system. This possibility would be in line with our pre-embedding EM result, with KCTD12b showing peak densities inside the active zone of ventral MHb terminals whereas KCTD8 shows peak expression in the peri-synaptic region. This could be an indication for the preferred interaction of Cav2.3 with KCTD12b over KCTD8 at rest.

To induce synaptic plasticity in the ventral MHb to IPN pathway, release of glutamate, activation of postsynaptic Ca^2+^-permeable AMPA receptors on GABAergic IPN neurons and subsequent retrograde release of GABA to activate presynaptic GABA_B_ receptors on MHb terminals is required (Koppensteiner et al., 2017). Therefore, KCTD-mediated modulation of both release probability and the persistence of GBR-mediated potentiation could determine a ventral MHb synapse’s propensity to undergo and express activity-dependent plasticity. If our hypothesis is correct, a synaptic terminal with a high KCTD8:12b ratio in the active zone would be expected to exhibit a high basal transmission at the cost of impaired potentiation after a strong stimulus. In contrast, a low KCTD8:12b ratio may be reflected by a low basal transmission but enhanced GBR-mediated potentiation after strong habenular activity. Futures studies may provide additional insights into whether physiological learning paradigms affecting the MHb-IPN pathway, such as the formation and extinction of aversive memories (Agetsuma et al., 2010; Koppensteiner et al., 2017; Melani et al., 2019; Zhang et al., 2016), could alter the ratio of KCTD8 to 12b.

Overall, our study provided new insights into the physiological role of presynaptic Cav2.3, GBRs and their auxiliary KCTD subunits in an evolutionary conserved neuronal circuit. Future studies will be required to identify the molecular mechanism underlying the GBR-mediated presynaptic potentiation on cholinergic MHb terminals. It remains to be determined whether the prominent presence of presynaptic KCTDs at other synapses (Müller et al., 2010) could be an indication of similar neuromodulatory function of KCTDs in different pathways of the brain.

## Materials and Methods

### Animals

Wild-type C57BL/6J (Jax, Bar Harbor, ME, USA; #000664) and BALB/cJ (Jax, #000651) mice were initially purchased from Jackson Laboratory. Homozygous KCTD KO lines were generated by the lab of Bernhard Bettler at the University of Basel (Schwenk et al., 2010). For KCTD8 KO line generation, exon 1 containing ATG and most of the open reading frame (ORF) of the Kctd8 gene (MGI:2443804) was replaced with a loxP-flanked neo. To generate KCTD12 KO mice, 5’ part of exon 1 containing complete ORF of the Kctd12 gene (MGI:2145823) was replaced with a loxP-flanked neo (Cathomas et al., 2015). Similarly, KCTD12b KO line was generated by replacing the 5’ part of exon 3 containing complete ORF of the Kctd12b gene (MGI:2444667) with a loxP-flanked neo. All KCTD KO lines had the neo removed by crossing founders with a Cre-deleter line, leaving one loxP site behind. The KCTD8/12d double KO line was generated by mating F2 hybrids of the parental lines. Background strains of KCTD KO lines were as follows: KCTD8 (C57BL/6J and 129), KCTD12 (C57BL/6J, 129, BALB/cJ), KCTD12b (BALB/cJ), KCTD8/12b (C57BL/6J, 129, BALB/cJ). To obtain Tachykinin1 (Tac1)-ChR2-EYFP mice, we crossed Tac1-Cre (Jax, #021877) with Ai32 (Jax, #012569) mice. All mice were bred at the preclinical facility of IST Austria on a 12:12 light-dark cycle with access to food and water *ad libitum*. All experiments were performed in accordance with the license approved by the Austrian Federal Ministry of Science and Research (Animal license number: BMWFW-66.018/0012-WF/V/3b/2016) and the Austrian and EU animal laws. Only male mice aged two to five months were used for all experiments.

### Transcardial perfusion for brain fixation

Mice were anaesthetized with a mixture of ketamine (90 mg/kg body weight) xylazine (4.5 mg/kg) solution intraperitoneally and 25 mM ice-cold phosphate buffer saline (PBS) was transcardially perfused through the left ventricle at a flow rate of 7 ml/min for 30 – 60 s. Subsequently, paraformaldehyde (PFA) solution was perfused for 12 minutes. PFA solutions of different concentrations were used for confocal imaging [4% PFA (TAAB) and 15% picric acid in 0.1 M phosphate buffer, PB], pre-embedding [4% PFA and 15% picric acid in 0.1 M PB + 0.05% glutaraldehyde(TAAB)] and SDS-digested freeze-fracture replica labeling (SDS-FRL, 2% PFA and 15% picric acid in 0.1 M PB). The pH of all PFA solutions was adjusted to 7.4 ± 0.05 with HCl. After perfusion, the brain was excised and placed in 0.1 M phosphate buffer (PB) and sectioned within 3 days. Slices of different thickness (50 µm for confocal microscopy and pre-embedding, 70 µm for SDS-FRL) were cut with a vibratome (Linear-Pro7, Dosaka, Japan) in ice-cold 0.1 M PB.

### Immunohistochemistry

Brain slices containing the IPN were washed with phosphate buffered saline (PBS) and subsequently incubated in blocking buffer (10% normal goat serum, 2% BSA, 0.5% Triton-X100 in 0.1 M PBS) for 1 h prior to incubation with primary antibodies: guinea pig anti-Cav2.3 [1 µg/ml, 2 overnight (O/N), Genovac], rabbit anti-GABAB1 [B17, 1 µg/µl, 1 O/N (Kulik et al., 2002)], rabbit anti-KCTD8 (1 µg/µl, Bettler lab, Univ. Basel), rabbit anti-KCTD12 (1 µg/µl, 1 O/N, Bettler lab, Univ. Basel), rabbit anti-KCTD12b [polyclonal, raised against a synthetic peptide comprised of the N-terminal amino acids 1-16 of KCTD12b; 1 µg/µl, 1 O/N, Bettler lab, Univ. Basel) (Metz et al., 2011; Schwenk et al., 2010). Brain sections were washed in PBS and subsequently incubated for 1 h in secondary antibody [1:500, Alexa-488 anti-guinea pig (Molecular Probes, Eugene, OR, USA) or Alexa-488 anti-rabbit (Molecular Probes)]. Sections were mounted onto glass slides and images were taken with an LSM 800 (Zeiss, Oberkochen, Germany) confocal microscope.

### Pre-embedding immunolabeling

Brain slices were washed in 0.1 M PB (2 times, 10 min each) and cryo-protected by incubation in 0.1 M PB containing 20% sucrose O/N at 4 °C. The next day, slices underwent three cycles of freeze-thawing by freezing the slices on liquid nitrogen for 1 min and thawing them in 20% sucrose on a hot plate (50°C) for 2 min. Slices were washed in 0.1 M PB (10 min) followed by washing in TBS (3 times, 20 min each). Free aldehydes were quenched by incubating slices in 50 mM glycine (Sigma Aldrich, St. Louis, MO, USA) in TBS (10 min). After washing in TBS (3 times, 20 min each), slices were blocked with blocking buffer (10% NGS, 2% BSA in TBS, 1 h). Primary antibody incubation was done with respective antibodies in 2% BSA solution for 2 O/N at 4 °C. The concentration of antibodies was 8 µg/ml for Cav2.3, 4 µg/ml for KCTD8, KCTD12 and KCTD12b and 2 µg/ml for GABA_B1_. Subsequently, slices were rinsed in TBS (3 times, 20 min each) and incubated in respective secondary antibodies (1:100) O/N at 4 °C in 2% BSA in TBS. For Cav2.3, 1.4 nm gold-conjugated anti-guinea pig antibody (Nanoprobes, Yaphank, NY, USA) and for all other antibodies, i.e. GABAB1, KCTD8, KCTD12 and KCTD12b, 1.4 nm gold-conjugated anti-rabbit antibody (Nanoprobes) was used. Slices were washed in TBS and PBS (2 times, 20 min each) followed by post-fixation in 1 % glutaraldehyde in PBS (10 min), washing in PBS (3 times, 10 min) and quenching of free glutaraldehyde in 50 mM glycine in PBS (10 min). Finally, slices were washed in PBS (3 times, 10 min each) and milli-Q (MQ) H_2_O (3 times, 5 min each).

For silver intensification of immunogold particles, Nanoprobes silver intensification (Nanoprobes) component A (initiator) and B (moderator) were mixed and vortexed, followed by the addition of component C (activator). After vortexing, slices were incubated in the mixture for 9 min 15 s in the dark. Tubes were tapped every 2 min for uniform intensification. Slices were washed with MQ water (3 times, 10 min each) and in 0.1 M PB (10 min), followed by post fixation in 1% OsO_4_ in 0.1 M PB (20 min in the dark). After osmification, slices were washed in 0.1 M PB (10 min) and in MQ water (3 times, 5 min each) and counter-stained in 1% uranyl acetate (Al-labortechnik, Zeillern, Germany) in MQ H_2_O (35 min in the dark). Subsequently, slices were serially dehydrated in ethanol solutions of different concentrations in ascending order up to 100% (50 – 95% ethanol in 5 steps, 5 min each; 100% ethanol 2 times, 10 min each) and then washed with propylene oxide (Sigma Aldrich; 2 times, 10 min each). Slices were then submerged in Durcupan resin (Sigma Aldrich; mixture of components A, B, C and D in proportion of 10:10:0.3:0.3 respectively) for 1 O/N at room temperature (RT).

For flat embedding, each slice was isolated on a silicon-coated glass slide, covered with an ACLAR® fluoropolymer film (Science Services, Munich, Germany) and incubated at 37 °C (1 h) followed by incubation at 60 °C (2 O/N). For re-embedding, tissue containing the rostral or lateral IPN was excised with a surgical blade, placed into the lid of a plastic tube (TAAB) which was then filled with Durcupan resin and incubated at 60 °C (2 O/N). Each resin block was trimmed using a Leica EM TRIM2 to remove the resin surrounding the sample. The resin above the sample in the trimmed block was further cut at 200 nm increments using a glass knife in the ultramicrotome Leica EM UC7 until the sample was exposed. 70 nm sections were cut with a diamond knife (Diatome Ultra 45 °). A small ribbon of floating sections was collected and mounted onto a copper-grid coated with formvar. Once the grid was dry, it was stored in a grid box for further observation in Tecnai10 (FEI; accelerating voltage 80 kV) or Tecnai 12 (FEI; accelerating voltage 120 kV) transmission electron microscopes.

### SDS-digested freeze-fracture replica preparation and labeling

Brain slices (70 µm) of mice transcardially perfused with 2% PFA in 0.1 M PB were prepared and the whole IPN was manually excised. Tissue was then incubated in 30% glycerol over night for cryo-protection. The next day, tissue samples were transferred into gold or copper carriers and frozen under high pressure (>300 bar) using an HPM010 (Leica, Wetzlar, Germany). Frozen samples were stored in liquid nitrogen until further processing. To craft freeze-fracture replicas, two gold carriers containing frozen tissue samples were placed in a carrier holder in liquid nitrogen, which was inserted into the freeze-fracture machine (BAF060, Leica) and left to equilibrate to −117 °C under high vacuum (2.0 × 10^−7^ – 1.0 × 10^−6^ mbar) for 20 min. Subsequently, tissue was fractured and a carbon layer (5 nm at a rate of 0.1 – 0.3 nm/s) was evaporated onto the tissue at 90°, followed by a platinum/carbon layer (2 nm at a rate of 0.06 – 0.1 nm/s) applied at 60° and another carbon layer (20 nm at a rate of 0.3 – 0.6 nm/s) applied at 90°. For the preparation of carbon-only replicas, the second layer consisted of a 5-nm carbon layer applied at 60°. After evaporation, replicas were removed from the machine and placed in tris-buffered saline (TBS, 50 mM). Subsequently, replicas were glued (tissue-side up) onto a finder grid and the glue (optical adhesive 61, Norland, Cranbury, NJ, USA) was hardened by applying UV light for 20 seconds and then transferred into SDS-solution containing: 2.5 % SDS, 20% sucrose in 15 mM Tris buffer (pH 8.3). Tissue was subsequently digested by incubating the replicas for 48 h at 60 °C under gentle agitation (50 rpm shaker), followed by incubation for 12 – 15 h at 37 °C under gentle agitation.

For immunolabeling of SDS-digested replicas, replicas were washed in washing buffer [containing: 0.1% Tween-20, 0.05% bovine serum albumin (BSA), 0.05% NaN_3_ pH=7.4] and incubated in blocking buffer (washing buffer + 5% BSA) for 1 h. Replicas were transferred to blocking buffer containing primary antibody (ginea pig anti-Cav2.3, 8 µg/µl) followed by incubation at 15 °C O/N. Thereafter, replicas were washed and incubated in blocking solution containing secondary antibodies (2-nm gold-conjugated anti-rabbit, 5-nm gold-conjugated anti-ginea pig or 10-nm gold-conjugated anti-rabbit, all diluted 1:30) at 15 °C O/N. The following day, the antibody labeling procedure was repeated for the next primary antibody. Labeled grid-glued replicas received a final carbon layer (20 nm) onto the labeled replica side using a High Vacuum Coater ACE600 (Leica), followed by dissolution of the glue in Dynasolve 711 (Dynaloy, Indianapolis, IN, USA) at 60 °C under gentle agitation (60 rpm) for 2 h. Solvent was subsequently removed by washing the grid in methanol (100, 95, 90, 70, 50%, 5 min each) and finally 100% ethanol. After short air drying, replicas were stored in grid boxes until observation under the transmission electron microscope.

### Acute brain slice electrophysiology

Mice were anesthetized with a mixture of ketamine (90 mg/kg) and xylazine (4.5 mg/kg) and transcardially perfused with ice-cold, oxygenated (95% O_2_, 5% CO_2_) artificial cerebrospinal fluid (ACSF) containing (in mM): 118 NaCl, 2.5 KCl, 1.5 MgSO_4_, 1 CaCl_2_, 1.25 NaH_2_PO_4_, 10 D-Glucose, 30 NaHCO_3_, (pH = 7.4). The brain was rapidly excised and coronal brain slices of 250 µm thickness were prepared with a Dosaka Linear-Pro7. For SNX-482 experiments in rostral IPN, brain slices were prepared at a 54° angle to allow for electrical stimulation of fasciculus retroflexus. Slices were recovered at 35 °C for 20 min and thereafter slowly cooled down to RT over the course of one hour. After recovery, one slice was transferred to the recording chamber (RC-26GLP, Warner Instruments, Holliston, MA, USA) and superfused with ACSF containing 2.5 mM CaCl_2_, 20 µM bicuculline methiodide, 50 µM hexamethonium-bromide and 5 µM mecamylamine hydrochloride at a rate of 3 – 4 ml/min at 32.0 ± 2.0 °C. Rostral or lateral IPN nuclei were visually identified using an infrared differential interference contrast video system in a BX51 microscope (Olympus, Tokyo, Japan). Electrical signals were acquired at 10 – 50 kHz and filtered at 2 kHz using an EPC 10 (HEKA, Lambrecht/Pfalz, Germany) amplifier. Glass pipettes (BF150-86-10, Sutter Instrument, Novato, CA, USA) with resistances of 3 – 4 MΩ were crafted using a P97 horizontal pipette puller (Sutter Instrument) and filled with internal solution containing (in mM): 130 K-Gluconate, 10 KCl, 5 MgCl_2_, 5 MgATP, 0.2 NaGTP, 0.5 EGTA, 5 HEPES; pH 7.4 adjusted with KOH. Whole-cell patch clamp recordings were performed in voltage-clamp mode at a holding potential of –60 mV and access resistance was constantly monitored via a –10 mV voltage step at the end of each sweep. Recordings with access resistances exceeding 20 MΩ or with changes in access resistance or holding current by more than 20% were discarded. To evoke glutamatergic excitatory postsynaptic currents (EPSCs) in rostral IPN neurons, voltage pulses (0.5 – 3.5 V, 0.2 ms duration) were applied with an ISO-Flex stimulus isolator (AMPI, Jerusalem, Israel) to a concentric bipolar stimulating electrode (CBBPC75, FHC, Bowdoin, ME, USA) located inside the IPN, ∼200 – 300 µm distal to the recorded neuron. For optogenetic stimulation in Tac1-ChR2 mice, blue light (λ = 465 nm) was emitted directly onto the lateral IPN through a 5 mm long mono fiber-optic cannula (fiber diameter 200 µm, total diameter 230 µm, Doric lenses, Quebec, Canada) connected to a PlexBright LED (Plexon, Dallas, TX, USA) with an optical patch cable (fiber diameter 200 µm, total diameter 230 µm, 0.48 NA). The LED was triggered via 200 – 290 mA current pulse (2 ms duration) from a LED Driver (LD-1, Plexon) which was controlled directly via the HEKA EPC10 amplifier. The sweep interval of all stimulation protocols (electrical and optogenetic) was 10 s. For the application of SNX-482 (1 µM, hellobio, Bristol, UK), 0.1% bovine serum albumin was added to the ACSF. For variance-mean analysis, ACSF with four different Ca^2+^ concentrations (1.5 – 8 mM) was applied to measure variance and mean EPSC amplitude at different release probabilities. Each Ca^2+^ concentration was washed in for 5 – 10 minutes. Once EPSC amplitudes stabilized, 15 – 20 consecutive EPSC responses were used for the calculation of mean EPSC amplitude and variance, followed by the wash-in of the next Ca^2+^ concentration. The order of application was 2.5 mM Ca^2+^, followed by 1.5 mM Ca^2+^, followed by 6 – 8 mM Ca^2+^ followed by 4 – 5 mM Ca^2+^. To measure the paired-pulse ratio (PPR) of two consecutively evoked EPSCs at 20 Hz, the PPRs of 20 – 30 EPSC pairs evoked at 10-second intervals were averaged. To study the effect of GBR activation on EPSC amplitude, R(+)-Baclofen hydrochloride (1 µM) was bath applied for three minutes and washed for 10 minutes. For PPR and variance-mean experiments, data from wild-type C57BL/6J and BALB/c mice were found to not differ significantly and thus were pooled.

### Cell culture transfection and electrophysiology

HEK293 cells, stably expressing the human Cav2.3 (R-type) channel α_1E-3_ splice variant (also called α1E-c; GenBank L29385), the Ca^2+^ channel auxiliary subunits human α_2b_δ-1 (M76559) and human β_3a_ (NM_000725) as well as the human potassium channel KCNJ4 (Kir2.3; U07364), were a kind gift from the laboratory of David J. Adams (University of Wollongong, Australia). Cells were maintained in culture as described before (Dai et al., 2008). For experiments, the Cav2.3-expressing HEK293 cells were transiently transfected with plasmids for the expression of Myc-tagged KCTD8 (AY615967) or KCTD12b (AL831725) using Lipofectamine 2000 (Invitrogen, Carlsbad, CA, USA) as described earlier (Schwenk et al., 2010). Plasmids for the expression of eGFP (Clontech, Kyoto, Japan) were co-transfected in order to identify cells expressing the plasmids. A total quantity of 0.75 µg of DNA was transfected in each well, and empty pCI plasmid was used to complete to this amount of total DNA. The control group contained only eGFP and empty vector. 2 **–** 4 days after transfection, cells were briefly subjected to Versene (ThermoFisher, Waltham, MA, USA) treatment for its dissociation and plated on glass coverslips and, at least 2 hours later, electrophysiology experiments were performed.

Whole-cell recordings were performed using fire-polished borosilicate patch pipettes (2 – 4 MΩ of tip resistance), filled with a cesium-based intracellular solution containing (mM): 110 CsMeSO_3_, 15 CsCl, 10 EGTA, 10 HEPES, 10 Tris-phosphocreatine, 0.1 NaGTP, 4 MgATP, pH 7.3 adjusted with CsOH. The extracellular solution contained (mM): 110 NaCl, 30 TEA-Cl, 10 HEPES, 10 D-glucose, 5 CsCl, 1 MgCl_2_ and 5 mM BaCl_2_, pH 7.4 adjusted with 40% TEA-OH. For patch-clamp experiments, isolated cells with a clear eGFP fluorescent signal were selected. The culture was maintained under continuous perfusion (∼1 ml/min) at room temperature. A MultiClamp 700B (Molecular Devices, San Jose, CA, USA) amplifier connected to a 1440A Digidata was used to acquire signals at 10 – 50 kHz, filtered at 2 – 8 kHz. After whole-cell formation, series resistance (<10 MΩ) was compensated by 85%. Linear leak currents and capacity transients were subtracted on-line using a −P/4 protocol. Current-voltage relationships were recorded from a holding potential of −80 mV using 25 ms depolarizations from −70 to 60 mV in 10-mV increments. Peak *I*_Ba_ was measured at each step and normalized to the capacitance of the cell to obtain the current densities, which were plotted as a function of the voltage step. I−V curves were fit with a standard Boltzmann equation as follows: *I*_Ba_ = *G*_max_(*V* − *V*_rev_)/(1+exp((*V*_0.5act_ − *V*)/*k*)), where *I*_Ba_ is the current measured at each test potential *V, G*_max_ is the maximal conductance, *V*_rev_ is the reversal potential, *V*_0.5act_ is the half-maximal voltage of activation and *k* is the slope factor. To study voltage-dependency of activation, a series of 25 ms voltage steps from −80 to 60 mV (with 10 mV increments) were applied, followed by a 10 ms step to 0 mV, where peaks of the evoked current tails were measured, normalized to the maximal peak and plotted against the corresponding voltage. To the study-steady state inactivation, a series of 2 seconds-long voltage steps from −120 to 10 mV (10 mV increments) was applied, followed by a test step to 10 mV (150 ms) where peaks of elicited currents were measured, normalized to the maximal response and plotted against the corresponding voltage. Both activation and inactivation curves were fitted using a Boltzmann sigmoidal, as follows: *I* = *I*_2_ *+* (*I*_1_ − *I*_2_) / (1+exp((*V*_0.5 act/inact_ – *V*_*t*_)/*k))*, where *V*_0.5 act/inact_ is the half-maximal activation or inactivation, and *k* is the slope factor.

### Co-Immunoprecipitation

Plasmids encoding N-terminally Flag-tagged KCTDs were described earlier (Seddik et al., 2012). Plasmids expressing C-terminally Flag-tagged Cav2.3 were obtained from GenScript. Cultured human embryonic kidney 293 (HEK293) cells were transfected using 2 μg/μl polyethylenimine (Sigma-Aldrich) with indicated plasmids for the expression of either Flag-KCTDs alone or in combination with Flag-Cav2.3. Cells were washed 48 h after transfection in ice cold PBS and lysed in NETN buffer (100 mM NaCl, 1 mM EDTA, 0.5% Nonidet P-40, 20 mM Tris-HCl, pH 7.4, supplemented with cOmplete EDTA-free protease inhibitor mixture (Roche, Basel, Switzerland)), followed by rotation for 10 min at 4 °C. Cell lysates were then cleared by centrifugation at 10 000 × g (4 °C, 10 min) and directly used for immunoblot analysis (Input) or for co-immunoprecipitation assay, in which they were precleared for 1 h using 30 μl (dry volume) of a 1:1 mixture of protein-A and protein-G-agarose beads (GE Healthcare, Chicago, IL, USA). Thereafter, lysates were incubated by rotating for 16 h at 4 °C in the presence of 2.5 μl of 0.3 μg/μl anti-Cav2.3 (CACNA1E) antibody (ACC-006, Alomone Labs, Jerusalem, Israel). The next day, 10 μl (dry volume) of a 1:1 mixture of protein-A and protein-G-agarose beads were added and incubated by rotating for 40 min at 4 °C. Afterward, they were washed with 5 × 1 ml of NETN buffer and pulled down proteins were eluted with 25 μl of 4 × sample loading buffer containing 200 mM DTT. Proteins were resolved using standard one-dimensional SDS-PAGE on 10% polyacrylamide gels (for 45 min at 70 mV, followed by an additional 1.5 h at 120 mV). For immunoblotting analysis, proteins were transferred to 0.45 μM polivinylidene fluoride membranes (Milipore, Burlington, MA, USA) for 2 h at 200 mA and probed with the primary antibodies rabbit anti-Flag (F7425, Sigma) and anti-Cav2.3 (ACC-006, Alomone Labs) in combination with peroxidase-coupled secondary donkey anti-rabbit antibodies (NA934, GE Healthcare, 1:10000).

### Data analysis and statistics

All statistical tests and graph preparations were done using Prism (GraphPad, San Diego, CA, USA) and figure assembly was done in Photoshop (Adobe, San Jose, CA, USA). To determine whether to use parametric or non-parametric statistical tests, Shapiro-Wilk test for normality of residuals was applied. Unless otherwise noted, averaged data is presented as mean ± SEM and P values < 0.05 were considered to indicate statistical significance. Power analysis (power = 0.8, α error = 0.05) was used to determine the appropriate sample size for SDS-FRL and variance-mean experiments using G*Power software (Univ. Kiel, Germany). Sample size in other experiments was based on those of similar experiments in previous studies. Masking was not used for any experiment. For the analysis of pre-embedding immunolabeled ultrathin sections, the presynaptic active zone was manually demarcated based on 1) rigid alignment of pre- and postsynaptic membranes and the presence of a postsynaptic density with the same length as the presynaptic active zone, as previously described (Rubio et al., 2017). Active zone length and silver-intensified gold particle densities and distances were measured using Reconstruct software. In SDS-FRL samples, demarcation of presynaptic active zones was performed manually based on two morphological criteria: there had to be a visible alteration in surface curvature in the P-face and/or a concentration in intramembrane particles (IMPs) of variable sizes that appeared clearly different from those of the surrounding P-face. For density measurements, incomplete active zones were analyzed whenever there was a visible transition to the postsynaptic E-face that displayed the characteristic IMP clusters of a glutamatergic postsynapse (Tanaka et al., 2005). For confirmation, the complete active zone area demarcated based on protoplasmic surface depression and concentrated IMP cluster were compared with the area of complete postsynaptic E-face IMP clusters. In addition, the area of manually demarcated complete active zones were also compared with those demarcated using a mixture of active zone-marker antibodies for RIM1/2 (5 µg/ml; Synaptic Systems, Göttingen, Germany), neurexin (5 µg/ml) (Miki et al., 2017), CAST (3 µg/ml) (Hagiwara et al., 2018). To analyze clustering of molecules (Cav2.3, GABA_B1_, KCTD8, and KCTD12b), we performed 100 Monte Carlo simulations using GPDQ software (Lujan et al., 2018). For each simulation, the same number of particles was redistributed randomly, with each pixel having the same probability of becoming the center of a particle, on the demarcated area of interest under the condition that two particles could not be closer to each other than 10 nm. We then compared nearest neighbor distances (NNDs) between real particles with the NNDs of simulated particles using Kolmogorov-Smirnov test.

## Acknowledgements

We are grateful to Akari Hagiwara and Toshihisa Ohtsuka for CAST antibody, and Masahiko Watanabe for neurexin antibody. We thank David Adams for kindly providing the stable Cav2.3 cell line. This project has received funding from the European Research Council (ERC) and European Commission (EC), under the European Union’s Horizon 2020 research and innovation programme (ERC grant agreement No. 694539 to Ryuichi Shigemoto and No. 692692 to Peter Jonas) and the Swiss National Science Foundation Grant 31003A-172881 to Bernhard Bettler.

